# Proton migration on biological membranes: Lipid phase, temperature, and composition dependence of proton transfer processes and membrane proton barrier

**DOI:** 10.1101/2024.10.11.617755

**Authors:** Ambili Ramanthrikkovil Variyam, Mateusz Rzycki, Ramesh Nandi, Alexei A. Stuchebrukhov, Dominik Drabik, Nadav Amdursky

**Affiliations:** Schulich Faculty of Chemistry, Technion – Israel Institute of Technology, Haifa, 3200003, Israel; Department of Biomedical Engineering, Wroclaw University of Science and Technology, Wroclaw, 50-370, Poland; Department of Chemistry, University of California at Davis, One Shields Avenue, Davis, California 95616, USA; Department of Chemistry, University of Sheffield, Sheffield S3 7HF, United Kingdom

**Keywords:** Membranes, Proton transport, Lateral diffusion, Membrane biophysics, Photoacids

## Abstract

Biological membranes play a major role in diffusing protons on their surfaces between transmembrane protein complexes. The retention of protons on the membrane’s surface is commonly described by a membrane-associated proton barrier that determines the efficiency of protons escaping from surface to bulk, which correlates with the proton diffusion (PD) dimensionality at the membrane’s surface. Here, we explore the role of the membrane’s biophysical properties and its ability to accept a proton from a light-triggered proton donor situated on the membrane’s surface and to support PD around the probe. By changing lipid composition and temperature, while going through the melting point of the membrane, we directly investigate the role of the membrane phase in PD. We show that the proton transfer process from the proton donor to the membrane is more efficient in the liquid phase of the membrane than in the gel phase, with very low calculated activation energies that are also dependent on the lipid composition of the membrane. We further show that the liquid phase of the membrane allows higher dimensionalities (close to 3) of PD around the probe, indicating lower membrane proton barriers. In the gel phase, we show that the dimensionality of PD is lower, in some cases reaching values closer to 1, thus implying specific pathways for PD, which results in a higher proton recombination rate with the membrane-tethered probe. Computational simulations indicate that the change in PD between the two phases can be correlated to the membrane’s ‘stiffness’ and ‘looseness’ at each phase.

**Significance statement:** Proton diffusion on the surface of biological membranes serves a vital role in migrating protons into bioenergetic systems. Here, we explore how the biophysical properties of the membrane determine proton migration and proton retention on the surface of the membrane, i.e., the membrane proton barrier. We show that the membrane phase, which is also influenced by lipid composition, has a crucial role in the proton circuity of biological membranes. We found that the gel phase reduces the proton diffusion dimensionality and that the proton barrier is determined by lipid composition. Our results highlight the complexity of proton migration on the surface of biological membranes and the associated biophysical parameters that influence the proton diffusion process.

## Introduction

Proton transfer (PT) reactions are at the heart of bioenergetic systems, where PT occurs between the two sides of a biological membrane via transmembrane protein complexes (1, 2). Much scientific attention was given to the question of how protons reach the transmembrane protein complexes, whether from the bulk aqueous medium or the surface of the membrane. Such studies resulted in our understanding that the surface of biological membranes can support lateral proton diffusion (PD) with an apparent proton barrier between the surface of the membrane and the bulk (3–8). This proton barrier can result in a delayed equilibrium between protons on the surface of the membrane and the ones in the medium surrounding it, which, in turn, can explain the observed lateral long-range PD on the surface of the membrane. Nevertheless, lateral long-range PD and the mechanism behind the proton barrier are not well understood. Also, the nature of the delayed equilibrium process is mysterious; while some suggest a quasi-equilibrium model between protons on the surface and bulk (9–11), i.e., fast desorption and adsorption of protons from the surface to the bulk, others suggest a non-equilibrium model (12–14), i.e., the retention of the protons on the surface of the membrane. While the initial studies targeting the membrane-related proton barrier proved the capability of biological membranes to support lateral PD (3–5, 15), more recent studies targeted the role of the membrane composition in this process (14, 16–22). A fundamental biophysical property of the membrane that was commonly overlooked in most studies concerning PD on the surface of biological membranes is the membrane phase, liquid (fluid) vs. gel phase. Each biological membrane is characterized by a transition melting temperature (T_m_), whereas above the T_m_, the membrane is in its liquid phase, and below it is in the gel (ordered) phase. In this study, we directly tackle the role of the membrane phase, which is also related to the membrane composition, on the ability of the membrane to support PD and the nature of the proton barrier. By changing the composition of the membrane and performing temperature-dependent measurements going through the T_m_, we can resolve the changes in the PD and other PT processes happening on the surface of the membrane and decipher the mechanism of PT.

To measure the PT reactions on the surface of the biological membrane, we use here our recently developed probe that can be tethered to the surface of the membrane and can release, i.e., inject, a proton to the surface of the membrane on demand following light excitation (19, 23). The probe is based on the pyranine photoacid to which long alkyl chains were attached, a probe that we term C_12_HPTS (**Figure 1**). Following excitation and owing to the low p*K_a_* value in the excited state of the probe, it undergoes a deprotonation process known as excited-state PT (ESPT):

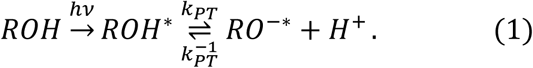

**Figure 1:**
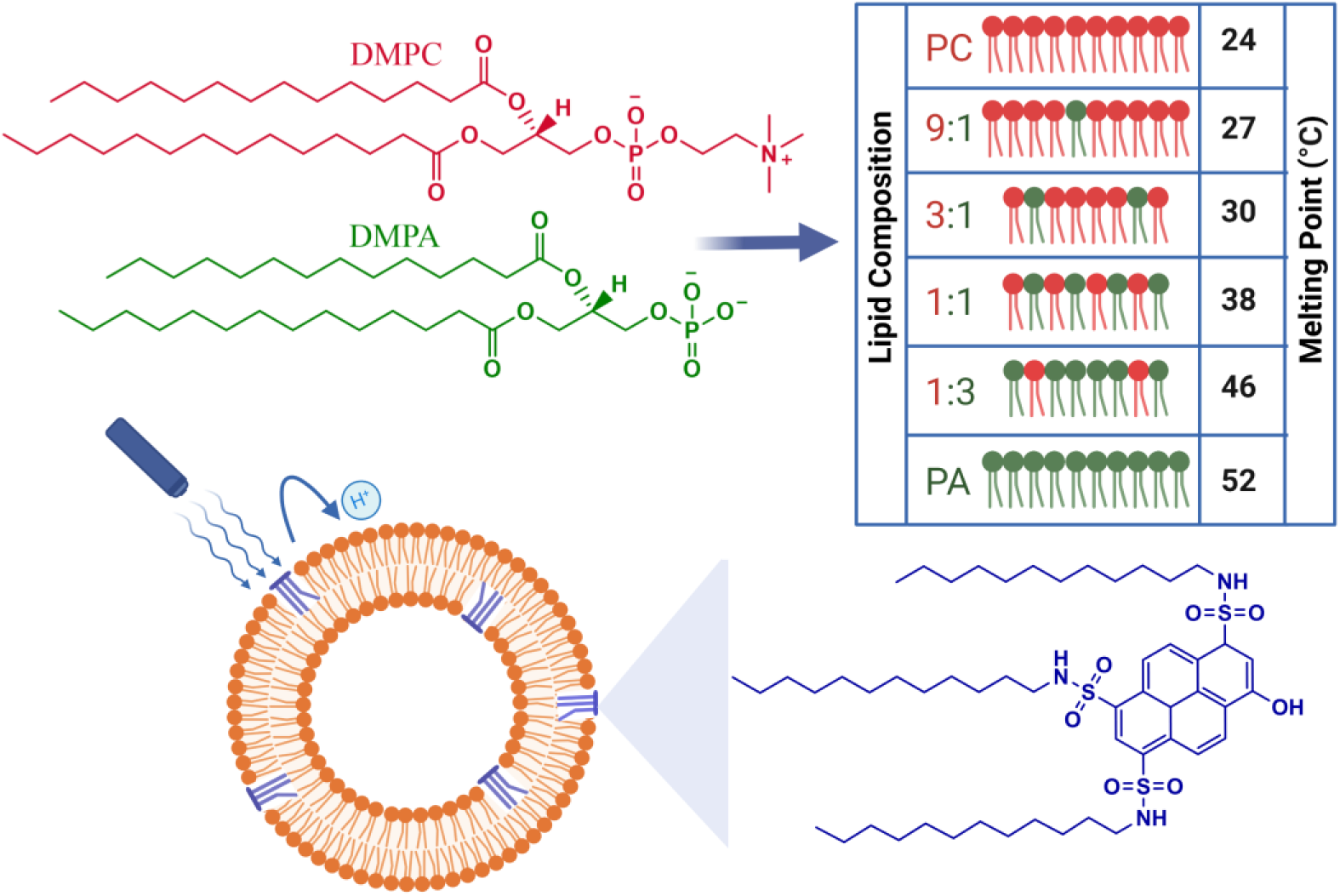
Molecular structures of DMPC and DMPA lipids and the ratios of these lipids used for making six different vesicles (top panel). Schematic diagram of light-induced PT from the membrane-tethered photoacid and the molecular structure of C_12_-HPTS (bottom panel).

Following deprotonation, the released proton to the surface of the membrane can either undergo lateral PD, escape to the bulk medium surrounding the membrane, or undergo geminate proton recombination with the deprotonated excited probe (*RO*^−∗^). In this way, our probe serves as both the proton donor, the *RO*^−^ form, and the proton acceptor, the *RO*^−∗^form. Since the protonated and deprotonated states of the probe emit at different wavelengths, we can detect the population of each state individually. Using steady-state and time-resolved fluorescence measurements, together with our recently developed model for the mentioned ESPT process (20), we can gain valuable information on the ESPT rate (*k_PT_*), the recombination rate (*k^−1^_PT_*), and the dimensionality of the PD process surrounding the probe. Since we observe only the excited-state of the probe in our measurements, and taking into account its few nanosecond lifetime, we can follow PD only at time scales of dozens of nanoseconds. Nonetheless, this time scale corresponds to PD lengths of >10 nm, which has physiological relevance as it is more than the common distance between protein complexes in bioenergetic systems.

## Results

### The membranes of this study

All the membranes we use in this study are small unilamellar vesicles (SUVs). Recently, we explored the role of membrane composition on the ESPT processes from C_12_HPTS and the related dimensionality of PD by using different SUVs, differentiated by the ratio of POPC to POPA (1-palmitoyl-2-oleoyl-glycero-3-phosphocholine and 1-palmitoyl-2-oleoyl-sn-glycero-3-phosphate, respectively) (21). Owing to the negatively charged POPA nature, it was expected to observe a faster ESPT process and a slower recombination rate (further discussion below) as more POPA lipids are within the membrane. However, we observed a complicated non-monotonous change in the ESPT parameters. We attributed the peculiar observation to a change in the membrane structure at low concentrations of PA within PC, which was shown to interfere with the PT capabilities of the membrane (21). In this study, we also use membranes with different ratios of PC:PA, but now we use the DMPC and DMPA (DM=1,2-dimyristoyl, i.e., 14:0 PC or PA). Unlike the POPC:POPA system, where all the different membrane compositions have their T_m_ below room temperature (RT), i.e., they are all in their liquid phase at RT, the DMPC:DMPA system has a large accessible range of T_m_ values between 24°C (for DMPC) and 52°C (for DMPA), hence, allowing us to follow the PT processes at different phases (**Figure 1** for the molecular structure of DMPC and DMPA, the ratios used in this study, and the T_m_ values of each membrane). Furthermore, while comparing the DM system to the PO system, we can compare the PT properties of a membrane with the same headgroup but at different phases at a given temperature.

In terms of the size of the vesicles used, all the different SUVs used in this study were of a similar size with a diameter of ∼100 nm, as estimated using dynamic light scattering (DLS) (**Figure S1**). The density of the inserted probe (1% vs. lipids) is such that the averaged distance between the probes is of the order of 7 nm, which is the same order of magnitude to the distance of protein proton channels/pumps within membranes. Another important system criterion is that the C_12_HPTS probe should be protonated (in its ground state) at the pH value of the solution used to allow the ESPT process in the excited-state. To verify it, we performed pH titration and followed the UV-vis absorption of the probe in each membrane composition used here (**Figure S2** and **Table S1** for the calculated apparent ground state pK_a_ values of the probe in the membranes).

### Steady-state fluorescence measurements

Steady-state fluorescence measurements can give us straightaway direct evidence that C_12_HPTS deprotonated by the emergence of a fluorescence peak (at ∼550 nm) associated with the deprotonated probe (*RO*^−∗^), whereas the peak of *RO*^−^is at ∼470 nm. **Figure 2** shows the normalized temperature-dependent steady-state fluorescence measurements at the temperature range of 10°-70°C for the DMPC and DMPA membranes and the different ratios between them used in this study (**Figure S3** for the graphs for POPC and POPA). As seen in the figure, the predominant peak at all measurements is the *RO*^−∗^ one (which is also undergoing a bathochromic shift as a function of temperature due to faster solvation at higher temperatures). However, the *RO*^−∗^/*RO*^−^is considerably changing as a function of temperature (**Figure 2**, insets, and **Table S2**). As can be observed in Eq.(1), the stationary (steady-state) population of *RO*^−∗^ is defined by the *k_PT_* and *k^−1^_PT_* rates as well as the deactivation rate (*k*_rx_) of the excited *RO*^−∗^, so as:

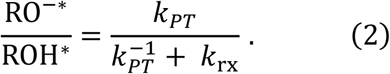

**Figure 2.**
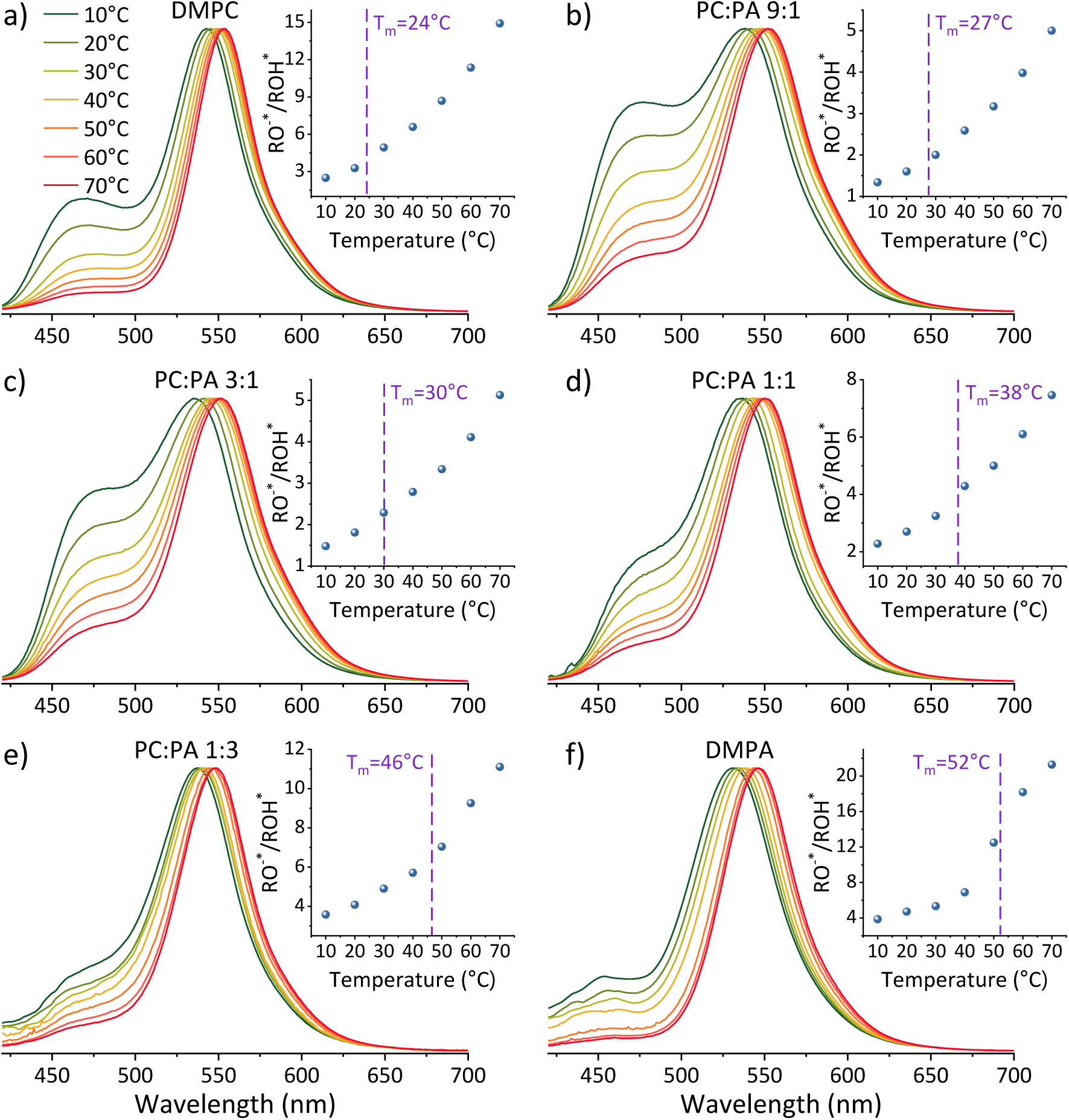
Normalized temperature-dependent steady-state fluorescence measurements of C_12_HPTS within membranes composed of a) DMPC, DMPC:DMPA ratios of b) 9:1, c) 3:1, d) 1:1, e) 1:3, and f) DMPA. The insets show the *RO*^−∗^/*RO*^−^as a function of temperature.

At this stage, we can already see a general trend where below the melting point, when the membrane is in its gel phase, the change in the *RO*^−∗^/*RO*^−^as a function of temperature is modest compared to the change in the ratio above the T_m_ when the membrane is in its liquid phase (the insets of **Figure 2**). However, from the steady-state fluorescence measurements alone, we cannot extract the needed PT rates to understand the PD process (as both the *k_PT_* and *k^−1^_PT_*rates are unknown and temperature-dependent), which leads us to the time-resolved measurements.

### Time-resolved fluorescence measurements

In line with Eq.(1), the time-resolved measurements can resolve between different processes happening in the excited-state of the probe at different times after the excitation process 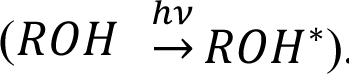. The first process is the ESPT process, the deprotonation of the excited probe, which is a fast process on the dozens to hundreds of picoseconds. This process reflects how good the surface of the membrane is in accepting the proton from the excited probe. A poor proton acceptor will result in a slow ESPT process, as we showed previously for cationic membranes (19), whereas a good proton acceptor will result in a fast ESPT, as we showed for anionic (POPA) membranes. Following the fast initial ESPT, there are two competing processes: the reverse proton recombination process (with *RO*^−∗^) for reforming *RO*^−^and the PD process, resulting in the escape of protons from *RO*^−∗^.

**Figure 3** shows the time-resolved decay of *RO*^−^for the DMPC and DMPA membranes and the different ratios between them used in this study (**Figure S4** for the graphs for POPC and POPA), measured at different temperatures (from 10°-70°C). Already in this stage, it is evident that the change in temperature changes both the initial fast process and the subsequent slow decay tail.

The fast initial ESPT process can be decoupled from the PD parameters of the accepting medium and the subsequent proton geminate recombination process, whereas the depletion in the excited ROH population (*p_ROH∗_*(*t*)) at very early times (dozens to hundreds of picoseconds) can be expressed as (20):

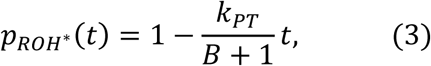

where B is a Boltzmann factor related to the charge of *RO*^−∗^ and corresponding proton attraction (20). As stated, the subsequent slower process is more complicated. As shown, both by us (20) and by the seminal work of Agmon and coworkers on photoacids (24–26), at the longer timescales of the process (tens of ns), the depletion of *p_ROH∗_*(*t*) decays as a power law, which is dependent on the dimensionality of PD around the probe (*d*):

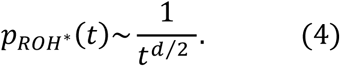

Accordingly, we can use our time-resolved measurements of all the different membranes used in this study at different temperatures (from 10°-70°C) to extract *k_PT_* from the first ∼100 ps of the decay, and *d* from later (>2 ns) in the decay (**Figure S5**, **Table S3**). The time-resolved measurements at the position of *RO*^−∗^ (**Figure S6**) can be used to calculate its *k*_rx_at the different membrane models and temperatures. By knowing, *k_PT_*, *k*_rx_, and *RO*^−∗^/*RO*^−^, we can extract the recombination rate *k^−1^_PT_*using eq. (2) (**Table S4**). It should be noted that in line with our previous study showing an anomalous change in *k_PT_* and *k^−1^_PT_*as a function of the %POPA in POPC vesicles at RT (21), we also observe the same anomaly for the DMPC:DMPA system at both the high-temperature liquid phase and the low-temperature gel phase (**Figure S7**). Nonetheless, our focus here is the change in the PT and PD properties as a function of temperature for different phases of a given membrane.

## Discussion

### Proton transfer and proton diffusion at different membrane phases and temperatures

**Figure 4** shows the parameters of *k_PT_*, *k^−1^_PT_*, and *d* (as in **Table S3** and **Table S4**) as a function of temperature for all the different membrane models along with the T_m_ of each membrane.

The figure highlights several important factors that are temperature-dependent and for which, the membrane phase dramatically changes them, which will be discussed below.

*The PT rate* (*k_PT_*) *– how well the membrane surface accepts a proton from the probe:* The rate *k_PT_* is directly estimated from the initial stages of the kinetics shown in **Figure 3**. In general, a PT process from a photoacid to the surrounding solution is a thermally active process, whereas the activation energy (E_a_, by fitting the calculated *k_PT_* to an Arrhenius equation) can also change as a function of temperature, resulting in higher values at lower temperatures (27, 28). The first important observation from our results is that the E_a_ of *k_PT_* is different between the liquid phase and the gel phase of a certain membrane. Furthermore, the change in the calculated E_a_ between the two phases of the membrane is related to the composition of the membrane. In DMPC membranes and membranes rich in DMPC (up to the ratio of 1:1 for DMPC:DMPA), the E_a_ calculated for the gel phase is higher than the one calculated for the liquid phase. However, in DMPA membranes and the ratio of up to 1:3 of DMPC:DMPA, the E_a_ calculated for the liquid phase was higher than that calculated for the gel phase. The latter trend was also observed for POPA membranes, having an accessible T_m_ value (28°C) (**Figure S8** comparing DMPA to POPA), hence highlighting the role of the headgroup in this observation. This finding is surprising since, as stated, the E_a_ is expected to increase with decreasing temperatures. Interestingly, the liquid phase of PC-rich membranes and the gel phase of PA-rich membranes both result in a very low E_a_ of the PT process in the order of only 11-14 meV. This small value indicates almost activation-less proton dissociation from the excited state, given k_B_T at room temperature of 25meV. It should be noted that all the extracted E_a_ values here are on the lower end of what was observed with the pyranine photoacid in solution (27), which highlights the different environment of the membrane surface for accepting a proton compared to the bulk medium.

**Figure 3.**
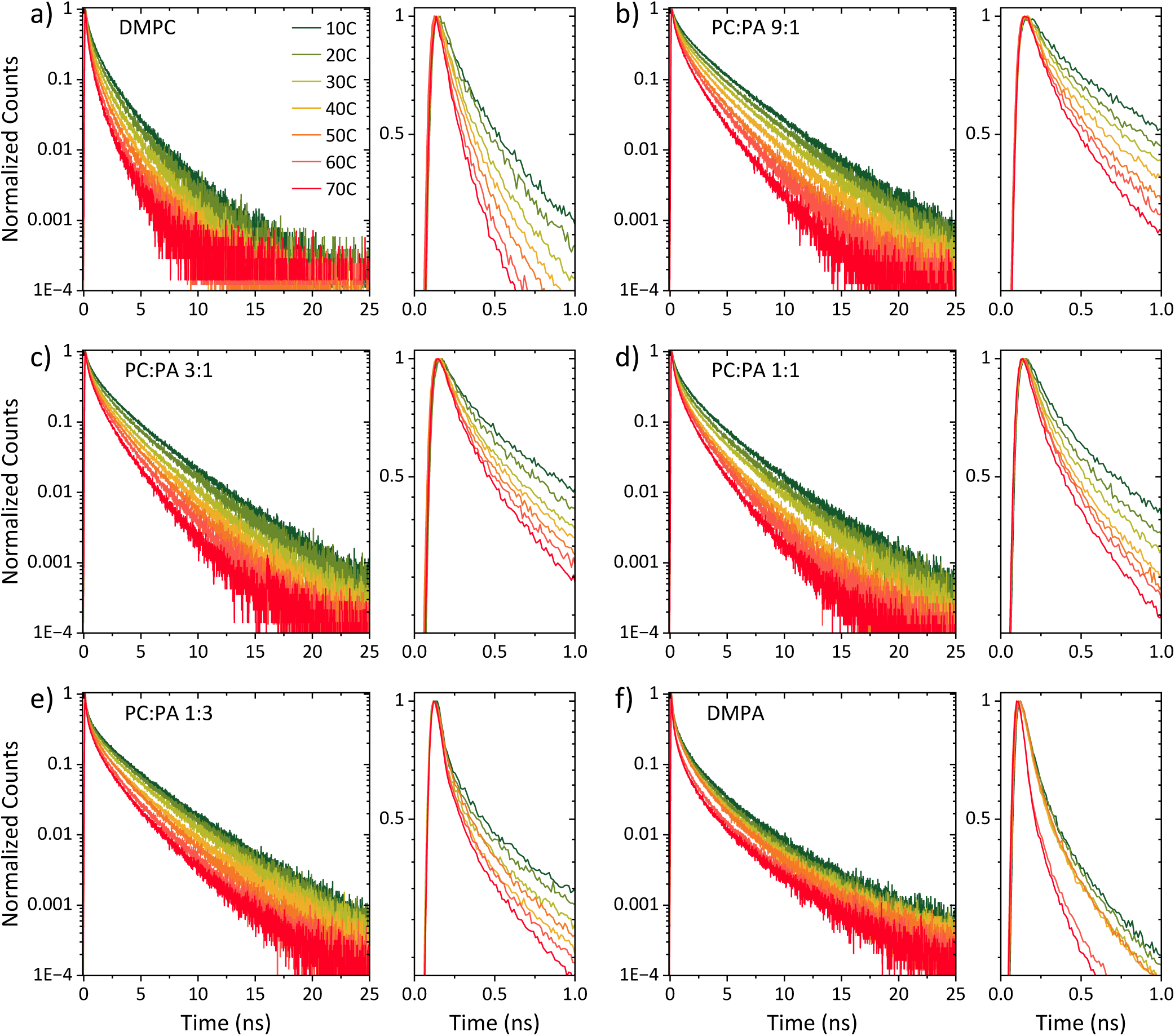
Normalized temperature-dependent time-resolved fluorescence measurements of C_12_HPTS within membranes composed of a) DMPC, DMPC:DMPA ratios of b) 9:1, c) 3:1, d) 1:1, e) 1:3, and f) DMPA. The right panels zoom in on the first nanosecond of the decay.

The magnitude of the proton dissociation rate *k_PT_*∼10^10^ s^-1^ that we obtain together with the magnitude of activation energy (enthalpy), E_a_ ∼10 meV, can be used by the transition state theory, *k_PT_* = 10^13^exp (−Δ*G*/*RT*), to calculate the activation entropy part (TΔS) of ΔG, which comes out to be around -180 meV. The latter estimate can be interpreted in terms of the fraction of thermal configuration states of the probe within the membrane with strong hydrogen bonds that lead to proton release, 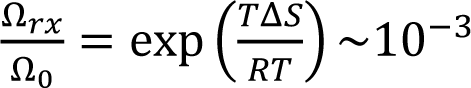. Such hydrogen bonds are transient, and the obtained number is a statistical fraction of total configurations that result in the reaction.

*The proton recombination rate* (*k^−1^_PT_*) *– how fast the membrane returns the proton back to RO*^−∗^: Given the above evaluations of *k_PT_*, and the estimate of relaxation rate *k*_rx_ (**Figure S6**), we can evaluate the rate *k^−1^_PT_* (using Eq. (2)). We found that this rate is in the order of a few 10^9^ s^-1^, depending on the composition of the membrane and temperature (**Figure 4**). When discussing proton recombination with the deprotonated photoacid in its excited state we need to differentiate between two sources of protons: 1) the geminate proton, i.e., the proton that was released from the photoacid to the surface of the membrane, 2) other protons from the aqueous environment surrounding the membrane and the probe. For investigating the contribution of protons coming from the medium, we supplemented our measurements at neutral pH (7.4) with measurements at low pH (3.6), meaning a change of 4 orders of magnitude in the proton concentration of the medium (**Figure S9**). In our measurements, we found that the change in *k^−1^_PT_* is only around two to three folds (depending on the temperature) higher at pH 3.6 (**Figure S9**) than at pH 7.4. Accordingly, we can safely claim that the main contribution to the magnitude of *k^−1^_PT_*is coming from geminate protons.

**Figure 4.**
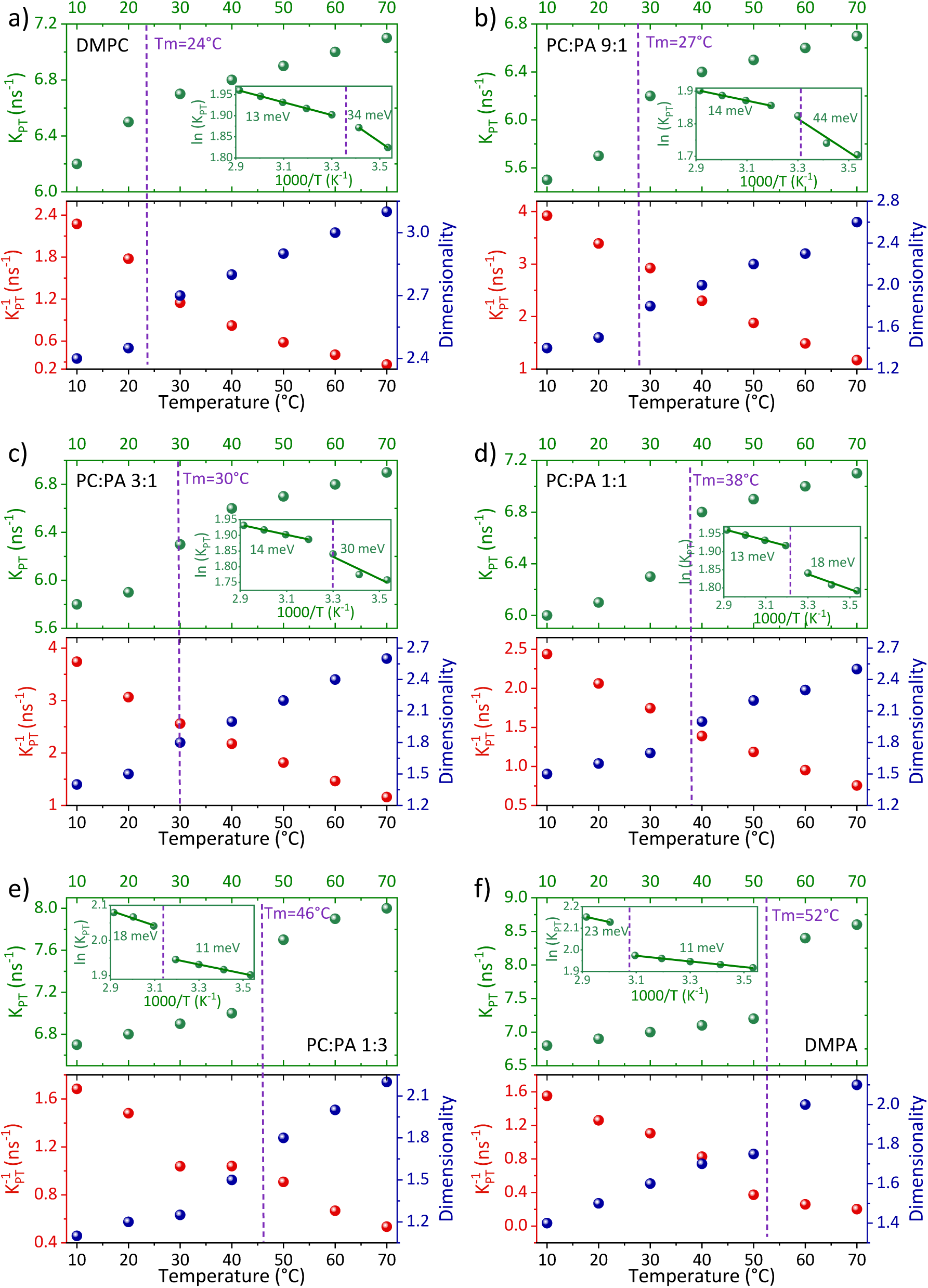
The extracted *k_PT_* values (top panels, green dots) from the initial decay and the extracted dimensionality (*d*) values (bottom panels, right axis, blue dots) from the slow tail of the decay of *RO*^−^ (see fitting in **Figure S5**), together with the extracted *k^−1^_PT_* values (bottom panels, left axis, red dots) using eq. (2) for membranes composed of a) DMPC, DMPC:DMPA ratios of b) 9:1, c) 3:1, d) 1:1, e) 1:3, and f) DMPA. The insets of the top panels are the same data plotted in an Arrhenius type (ln(*k_PT_*) vs. 1000/T) together with the calculated activation energy at each membrane phase.

Moreover, if we assume that the recombination occurs with the typical bimolecular rate of 10^11^ M^-1^s^-1^, the concentration corresponding to the geminate proton around the probe can be evaluated from the relation *k^−1^_PT_*=10^11^[H^+^]_g_. For a rate in the magnitude of *k^−1^_PT_*∼10^9^ s^-1^, we find that the geminate proton concentration is [H^+^]_g_=10^-2^ M, i.e. the effective geminate is at pH=2, i.e. much lower than the pH of biological system. Interestingly, this geminate concentration corresponds to one proton per ∼100 nm^-3^, which means the geminate proton explores the region of some 4-5 nm around the probe before recombining with the probe. This distance is also in the same order of the calculated distance between the probes on the surface of the membrane.

As for the change in the calculated *k^−1^_PT_* values, we observed a gradual decrease in *k^−1^_PT_* as a function of temperature, without a major indication of a different rate of change between the liquid and gel phases of the membrane. This means that the recombination of the proton coming from the membrane with *RO*^−∗^ is more efficient at low temperatures, which is in contrast to the discussed *k_PT_*. Moreover, the change in the magnitude of *k^−1^_PT_*is also larger than the discussed change in *k_PT_*, whereas for the DMPC/DMPA system and the different ratios of them, the change in *k^−1^_PT_* is several folds going from 10° to 70°C. Interestingly, the change in *k^−1^_PT_* as a function of temperature is lower for the PO systems than the DM ones, and especially while comparing POPC and DMPC (**Figure S10**). *What does it all mean?* Unlike *k_PT_*, which is not associated with the subsequent PD happening after proton dissociation, *k^−1^_PT_* is highly related to the PD parameters of the dissociated proton. If the dissociated proton can rapidly diffuse away from *RO*^−∗^, it will result in low *k^−1^_PT_* values, and vice-versa. Accordingly, the high *k^−1^_PT_* calculated for the low temperatures is indicative of the poor escape of protons at such temperatures, which is reasonable. The magnitude of the change in *k^−1^_PT_* as a function of temperature is then directly related to the capability of the membrane to enable lateral PD from the probe or the escape of protons from the membrane surface to bulk. We can define two important parameters in this context: the proton diffusion coefficient of the process and its dimensionality. Unlike previous models (26), our model for the ESPT process does not estimate the proton diffusion coefficient of the dissociated proton. However, as discussed, we can directly estimate the dimensionality of PD from the long-time component of the decay.

*The dimensionality (d) – how the geminate proton diffuses around the probe following dissociation*: The last parameter that is shown in **Figure 4** and will be discussed in this section is the dimensionality of the diffusion process of the geminate proton around the probe (20). We will start with DMPC and DMPA at the high-temperature regime, meaning they are both in their liquid state. In this condition, DMPA and DMPC show the dimensionality of 2 and 3, respectively, which is also in line with what was observed with POPA and POPC (21). These values mean that protons can diffuse laterally on the surface of PA membranes, while on the surface of PC membranes, they can also diffuse into the bulk medium above the membrane. This observation already indicates a lower proton barrier on the surface of PC membranes than on the surface of PA membranes, which is reasonable considering the negatively charged PA surface. At the high-temperature regime, the dimensionality extracted for the different DMPA:DMPC ratios falls between 2 and 3, whereas the higher the PA fraction, the closer it is to 2. The next important observation is that the extracted *d* values change as a function of temperature, whereas the lower the temperature, the lower the *d* value. Moreover, the magnitude of the change of *d* values as a function of temperature is generally smaller at the liquid phase of the membrane (at high temperatures) compared to a larger change in the gel phase (at low temperatures). For DMPC, the *d* values go down to 2.4 at 10°C. However, upon adding PA, and regardless of the amount of PA (even at the small DMPC:DMPA ratio of 9:1), the calculated dimensionality is significantly reduced, reaching values of ≤1.5 at the low-temperature gel phase of such membranes. Such low dimensionality values suggest a further restriction of the PD process from lateral dimensionality to lower ones, such as the previously discussed pathways/wires for protons on the surface of membranes (12, 29), and an increase in the proton barrier of the membrane.

To summarize this part of the discussion, our temperature-dependence studies of the different membranes reveal a clear distinction between the capability of membranes to accept a proton and support PD in their liquid vs. gel phase, which is also dependent on their composition. The activation energy for the PT process from the probe to the surface is generally low (<40 meV), whereas in membranes with high PC content, the E_a_ is higher in the gel phase, and in membranes with high PA content, the E_a_ is higher in the liquid phase. Following dissociation, the recombination rate of the proton with the deprotonated probe decreases as a function of increasing temperature due to the escape and diffusion of protons from the probe. The PD dimensionality is also temperature-dependent, showing larger values in the liquid phase than in the gel phase. We also observed that having PA in the membrane results in a lower dimensionality (<2) at the gel phases of such membranes.

### Computational analysis of the membrane’s biophysical and structural properties

The main remaining question is what biophysical and structural properties of the membrane that are changing as a function of temperature can explain our observations of the PT to the membrane surface and PD at the different phases of the membrane. To answer this question, we calculated a set of membrane-related parameters (see further details in the experimental section) for the different DMPC:DMPA ratios used here and at different temperatures covering the two sides of the T_m_: membrane thickness, area per lipid, lipid diffusion, and membrane bending (**Figure 5**).

**Figure 5.**
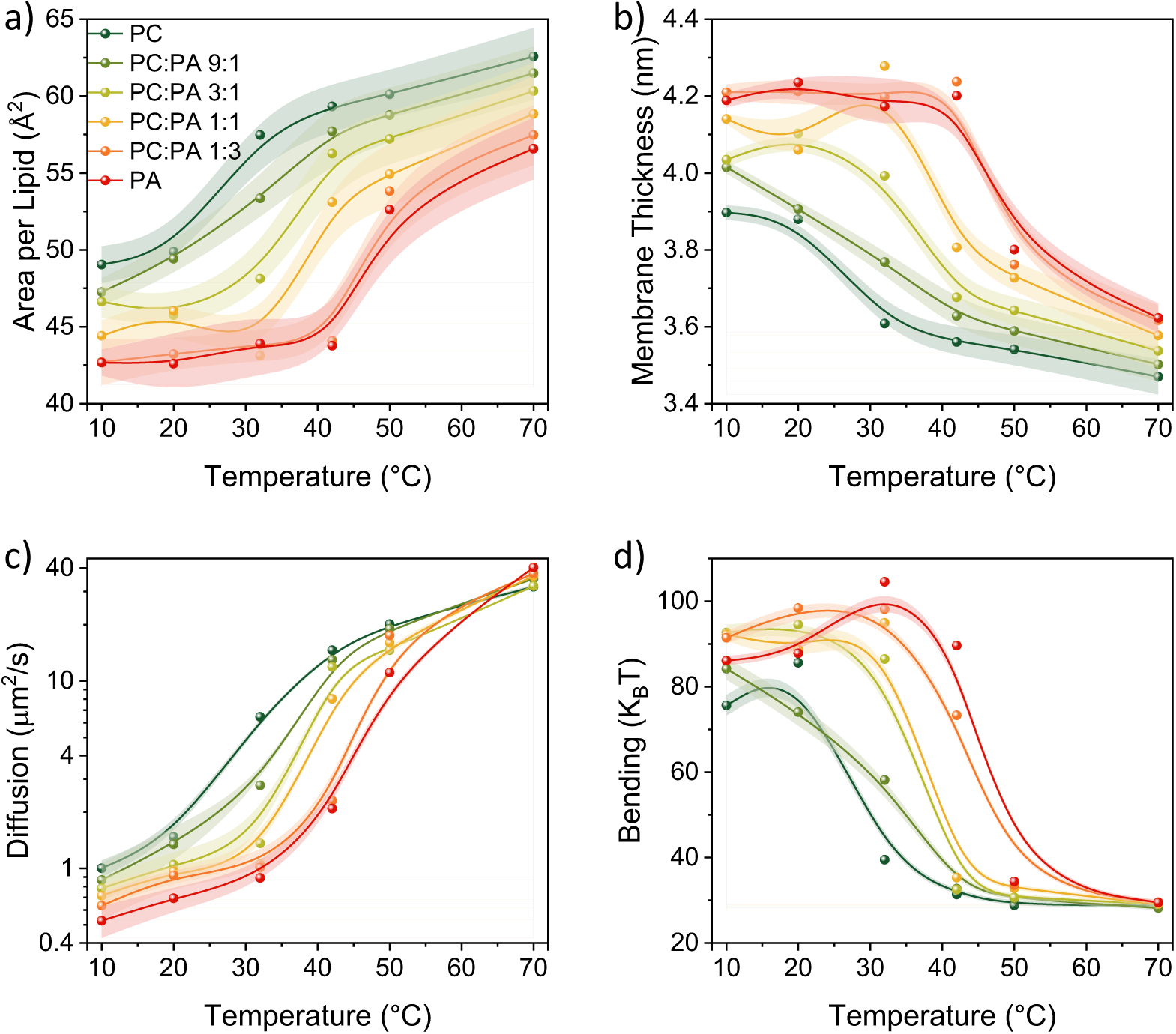
The calculated parameters of (a) area per lipid, (b) membrane thickness, (c) lipid diffusion, and (d) bending energy as a function of temperature for the different membranes used in this study composed of DMPA and DMPC.

The first notable observation from **Figure 5** is that all the biophysical properties simulated are experiencing a sharp change in their values around the T_m_ of the membrane composition, which also corresponds to what we found with the *k_PT_*values (**Figure 4**). According to our calculations, the membranes in their gel phase have a lower area per lipid, slower lipid diffusion, they are thicker, and with a higher bending energy than the membranes in their liquid phase. Hence, the efficient *k_PT_* values we observed in the liquid phase are associated with a ‘looseness’ structure of the membrane (high area per lipid, fast diffusion of lipids, low thickness, and low bending energy). This membrane configuration also allows higher dimensionality of PD, while for membranes with high content of DMPC, it also lowers the membrane proton barrier, i.e., the escape of protons from the surface. On the other side, the ‘stiffer’ membrane conformation at low temperatures supports lower dimensionality of PD, i.e., the emergence of proton pathways, which in turn increases the *k^−1^_PT_*values. Such ‘stiffer’ membrane conformation at low temperatures also changes the membrane configuration, as can be observed by the high bending (**Figure 5d**), which also results in high membrane thickness (**Figure 5b**), both due to the straightening of lipid tails.

Another notable observation from the computational analysis is that for membranes with a high content of DMPA (the red and dark orange curves, correspond to DMPA and PC:PA of 1:3, respectively, in **Figure 5**), the calculated parameters do not vary much as a function of temperature but only within the temperatures of their gel phase (primarily the area per lipid and membrane thickness). This observation might be correlated to the very low activation energy we found for the *k_PT_* values in this temperature range for the different membrane compositions.

### Conclusions

In summary, we focused in this study on the role of the membrane phase in several parameters of the membrane related to PT and PD: The PT from the membrane-tethered C_12_-HPTS photoacid to the surface of the membrane (as noted by *k_PT_*), the reverse process of proton recombination with the deprotonated photoacid in the excited-state (as noted by *k^−1^_PT_*), and the dimensionality of PD following the release of the proton from the photoacid. The latter property can also serve as an indicator of a change in the proton barrier of the membrane; dimensionality values closer to 3 are indicative of a low proton barrier and the escape of protons from the surface to the bulk. On the other extreme, low dimensionality values closer to 1 are indicative of a formation of proton pathways on the surface of the membrane, which is more restrictive than lateral PD on the surface of the membrane. By following the steady-state and time-resolved fluorescence of the photoacid as a function of lipid composition and temperature, going across the T_m_ of the membrane, we could reveal several important aspects of the PT and PD properties of biological membranes:

1. As expected, the PT rate from the photoacid to the membrane increases as a function of temperature, and we found the largest change while going from the gel to the liquid phase.
2. The activation energy of the latter PT process is very low (∼10-40 meV), and it is also dependent on the lipid composition. In membranes with a high content of DMPA, the activation energy is lower in the gel phase than in the liquid phase, while in membranes with a high content of DMPC it is the opposite, the activation energy is lower in the liquid phase.
3. The extracted back recombination process is also temperature-dependent, but unlike the PT process to the membrane, the rate of the reverse process (*k^−1^_PT_*) is decreasing as a function of temperature, i.e., it is higher in the gel phase, with no major ‘jump’ around the T_m_. The extracted kinetic parameters also indicate that the concentration of protons on the surface of the membrane following the photoacid deprotonation is orders of magnitude larger than the one in the solution, thus further implying the existence of a proton barrier.
4. The extracted dimensionality is also temperature-dependent, and it is increasing as a function of temperature. As one can expect, increasing the temperature can overcome the proton barrier of the membrane. Indeed, we observed that at high temperatures, and especially for membranes containing primarily DMPC, the extracted dimensionality is approaching 3, i.e., escape of protons from the surface to the bulk. On the other extreme, we found that except for pure DMPC membranes, at low temperatures, in the gel phase of the membrane, the dimensionality is reduced to below 2. This interesting observation implies the formation of proton pathways in the gel phase that restrict proton diffusion dimensionality, assisted by the presence of DMPA lipids.

In our study, we also performed temperature-dependent computational studies of several of the membrane’s biophysical properties: membrane thickness, area per lipid, lipid diffusion, and membrane bending energy, for the different lipid compositions used in this study. The computational result also shows a major change in each of the parameters around the T_m_ of the membrane. Such results imply a correlation between the observed PT and PD properties of the membrane to the other biophysical properties and suggest that the PT/PD properties observed for the gel phase might be due to the ‘stiffness’ of the membrane in that phase, while for the liquid phase, it might be due to the ‘looseness’ of the membrane in that phase.

## Materials and Methods

### Preparation of small unilamellar vesicles (SUVs)

All the lipids were purchased from Avanti Polar Lipids and used without further purification. A lipid solution of 2 mM was prepared in chloroform at a ratio of 1:100 for the probe to lipid molecules. The solvent was evaporated to and a dried lipid film was formed. Later, the lipid film was rehydrated with a 5 mM phosphate buffer of pH 7.4. Finally, the solution was extruded through a polycarbonate membrane (purchased from T&T Scientific) to obtain a homogenous solution. When needed, the pH was adjusted using 0.1 M NaOH or HCl solutions.

### Steady state spectroscopic measurements

Steady state UV-Visible absorption and fluorescence emission experiments were carried out using Agilent Cary 60 spectrometer and FS5 spectrofluorometer (Edinburg Instruments) respectively. Dynamic light scattering (DLS) experiments were measured using Zetasizer Nano ZS (Malvern Instruments). Differential scanning calorimetry (DSC) measurements were carried out using DSC-1 model from Mettler Toledo to find out the melting point of different lipid membranes.

### Time-correlated single photon counting experiments

Fluorescence life time of probe inside different vesicles were measured using a CHIMERA spectrometer (Light Conversion) with an excitation pulse at 400 nm where the time-resolved spectrum was collected using a hybrid detector (Becker & Hickl, HPM-100-07). The laser system includes a 10 W and 1030 nm Yb based PHAROS (Light Conversion) laser with a pulse of 190 fs, operating at 1 MHz with a maximum pulse energy of 10 μJ. The laser beam was seeded into an optical amplifier ORPHEUS (Light Conversion) for harmonic generation.

### MD simulations

The full description of the MD simulations is in the Supporting Information.

### Author contributions

A.R.V. and N.A. designed the research, A.R.V., M.R., R.N., and D.D. performed the research, A.A.S. contributed analytic tools, A.R.V., M.R., A.A.S., D.D., and N.A. analyzed the data and wrote the paper.

## Supporting information

Supporting Figures and Supporting Computational Information

## Acknowledgements

N.A. and A.A.S thank the United States - Israel Binational Science Foundation (grant number: 2018239 (NA) and A19-3374 (AAS)) for financial support. N.A. thanks the Lower Saxony – Israel Research Cooperation (grant number ZN3625) for financial support. M.R. thanks the National Science Centre, Poland (grant number 2022/45/N/NZ9/02130) for financial support and Wroclaw Centre for Networking and Supercomputing (http://www.wcss.pl) for providing computing resources (grant number 274).

## Declaration of Interests

The authors declare no competing interests.

## Notes

### Competing Interest Statement

The authors have declared no competing interest.

