## Supporting Figures and Supporting Computational Information for "Proton migration on biological membranes: Lipid phase, temperature, and composition dependence of proton transfer processes and membrane proton barrier"

**Table S1:** The ground state and excited state pKa of $C_{12}$-HPTS inside all the lipid mixtures with the absorption and emission maxima for ROH and $RO^-$ states.

| Liposome | Absorption (nm) | | Emission (nm) | | $pK_a$ | $\Delta pK_a$ | $pK_a^*$ |
|---|---|---|---|---|---|---|---|
| | $\lambda_{ROH}$ | $\lambda_{RO-}$ | $\lambda_{ROH}$ | $\lambda_{RO-}$ | | | |
| PC | 423 | 501 | 471 | 547 | 8.13 | 6.99 | 1.14 |
| PA:PC 1:9 | 420 | 503 | 471 | 544 | 9.34 | 7.15 | 2.19 |
| PA:PC 1:3 | 422 | 502 | 474 | 540 | 9.65 | 6.70 | 2.95 |
| PA:PC 1:1 | 418 | 501 | 474 | 540 | 10.13 | 6.90 | 3.23 |
| PA:PC 3:1 | 413 | 494 | 465 | 541 | 11.35 | 7.37 | 3.98 |
| PA | 412 | 486 | 460 | 541 | 11.83 | 7.33 | 4.50 |

**Table S2:** The ratio of $RO^{-*}$ to ROH* at different temperatures for all the membranes.

| Liposome | $RO^{-*}$/ROH* | | | | | | |
|---|---|---|---|---|---|---|---|
| | **10**°C | **20**°C | **30**°C | **40**°C | **50**°C | **60**°C | **70**°C |
| PC | 2.49 | 3.27 | 4.93 | 6.58 | 8.69 | 11.36 | 14.92 |
| PA:PC 1:9 | 1.34 | 1.60 | 2.00 | 2.59 | 3.17 | 3.98 | 5.00 |
| PA:PC 1:3 | 1.48 | 1.81 | 2.29 | 2.79 | 3.34 | 4.11 | 5.13 |
| PA:PC 1:1 | 2.28 | 2.70 | 3.25 | 4.29 | 5.00 | 6.10 | 7.46 |
| PA:PC 3:1 | 3.58 | 4.08 | 4.90 | 5.71 | 7.04 | 9.26 | 11.11 |
| PA | 3.87 | 4.72 | 5.35 | 6.90 | 12.50 | 18.18 | 21.28 |

**Table S3:** $k_{PT}$ (extracted from short time kinetics), B, d, and $\tau_0$ (extracted from long time kinetics) as a function of temperature for all the mixtures.

| **Temperature (°C)** | **Short time kinetics** | **Long time kinetics** | | |
|---|---|---|---|---|
| | $k_{PT}$ (ns$^{-1}$) | B | $d$ | $\tau_0$ (ns) |
| PC | | | | |
| 10 | 6.2 | 2.5 | 2.4 | 1.8 |
| 20 | 6.5 | 2.2 | 2.45 | 1.5 |
| 30 | 6.7 | 1.9 | 2.7 | 0.9 |
| 40 | 6.8 | 1.7 | 2.8 | 0.7 |
| 50 | 6.9 | 1.6 | 2.9 | 0.5 |
| 60 | 7.0 | 1.5 | 3.0 | 0.4 |
| 70 | 7.1 | 1.45 | 3.1 | 0.3 |
| PC:PA 9:1 | | | | |
| 10 | 5.5 | 2.8 | 1.4 | 1.8 |
| 20 | 5.7 | 2.7 | 1.5 | 1.4 |
| 30 | 6.2 | 2.35 | 1.8 | 1.0 |
| 40 | 6.4 | 2.2 | 2.0 | 0.9 |
| 50 | 6.5 | 2 | 2.2 | 0.8 |
| 60 | 6.6 | 1.9 | 2.3 | 0.7 |
| 70 | 6.7 | 1.8 | 2.6 | 0.5 |
| PC:PA 3:1 | | | | |
| 10 | 5.8 | 2.7 | 1.4 | 1.7 |
| 20 | 5.9 | 2.6 | 1.5 | 1.6 |
| 30 | 6.3 | 2.5 | 1.8 | 1.5 |
| 40 | 6.6 | 2.3 | 2.0 | 1.2 |
| 50 | 6.7 | 2.2 | 2.2 | 1.1 |
| 60 | 6.8 | 2.1 | 2.4 | 0.9 |
| 70 | 6.9 | 2.0 | 2.6 | 0.8 |
| PC:PA 1:1 | | | | |
| 10 | 6.0 | 2.5 | 1.5 | 1.5 |

| | | | | |
|---|---|---|---|---|
| 20 | 6.1 | 2.4 | 1.6 | 1.4 |
| 30 | 6.3 | 2.1 | 1.7 | 1.3 |
| 40 | 6.8 | 1.9 | 2.0 | 1.0 |
| 50 | 6.9 | 1.7 | 2.2 | 0.8 |
| 60 | 7.0 | 1.65 | 2.3 | 0.7 |
| 70 | 7.1 | 1.6 | 2.5 | 0.5 |
| PC:PA 1:3 | | | | |
| 10 | 6.7 | 2.5 | 1.1 | 1.4 |
| 20 | 6.8 | 2.3 | 1.2 | 1.3 |
| 30 | 6.9 | 2.1 | 1.25 | 1.25 |
| 40 | 7.0 | 1.95 | 1.5 | 1.1 |
| 50 | 7.7 | 1.8 | 1.8 | 1.0 |
| 60 | 7.9 | 1.7 | 2.0 | 0.9 |
| 70 | 8.0 | 1.65 | 2.2 | 0.8 |
| PA | | | | |
| 10 | 6.8 | 1.9 | 1.4 | 1.4 |
| 20 | 6.9 | 1.7 | 1.5 | 1.3 |
| 30 | 7.0 | 1.55 | 1.6 | 1.2 |
| 40 | 7.1 | 1.5 | 1.7 | 0.9 |
| 50 | 7.2 | 1.4 | 1.75 | 0.5 |
| 60 | 8.4 | 1.2 | 2.0 | 0.3 |
| 70 | 8.6 | 1.1 | 2.1 | 0.2 |

**Table S4:** $k_{PT}^{-1}$ calculated for all the mixtures at different temperatures.

| **Liposome** | $K_{PT}^{-1}$ | | | | | | |
|---|---|---|---|---|---|---|---|
| | **10**°C | **20**°C | **30**°C | **40**°C | **50**°C | **60**°C | **70**°C |
| PC | 2.27 | 1.78 | 1.15 | 0.82 | 0.58 | 0.40 | 0.26 |
| PA:PC 1:9 | 3.92 | 3.39 | 2.92 | 2.30 | 1.88 | 1.49 | 1.17 |
| PA:PC 1:3 | 3.74 | 3.06 | 2.56 | 2.18 | 1.82 | 1.47 | 1.16 |
| PA:PC 1:1 | 2.44 | 2.06 | 1.74 | 1.39 | 1.18 | 0.95 | 0.76 |
| PA:PC 3:1 | 1.68 | 1.48 | 1.04 | 1.04 | 0.91 | 0.67 | 0.53 |
| PA | 1.55 | 1.26 | 1.11 | 0.83 | 0.37 | 0.26 | 0.20 |

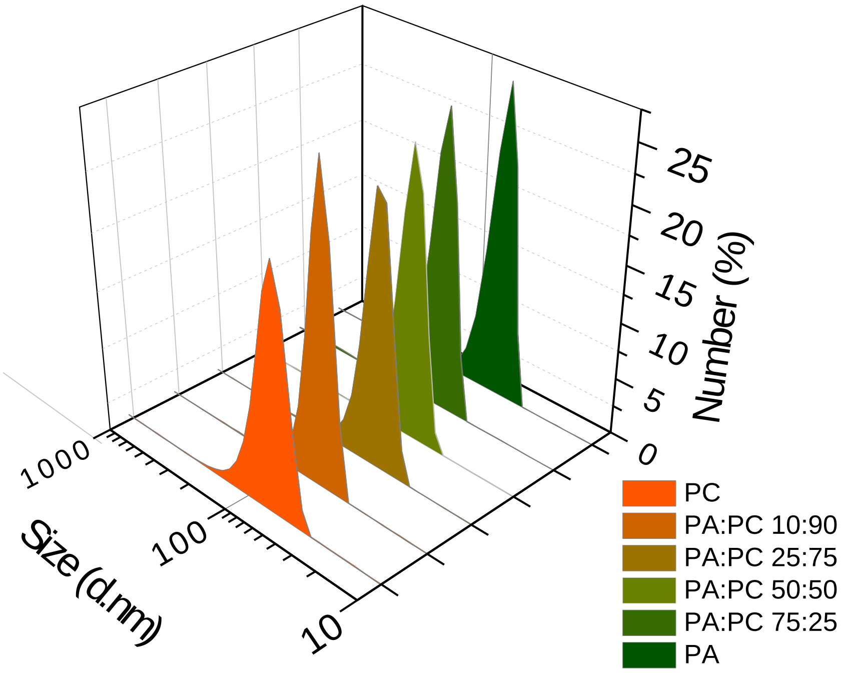


**Figure S1:** DLS measurements of all the mixtures showing the formation of monodispersed solutions of SUVs after extrusion.

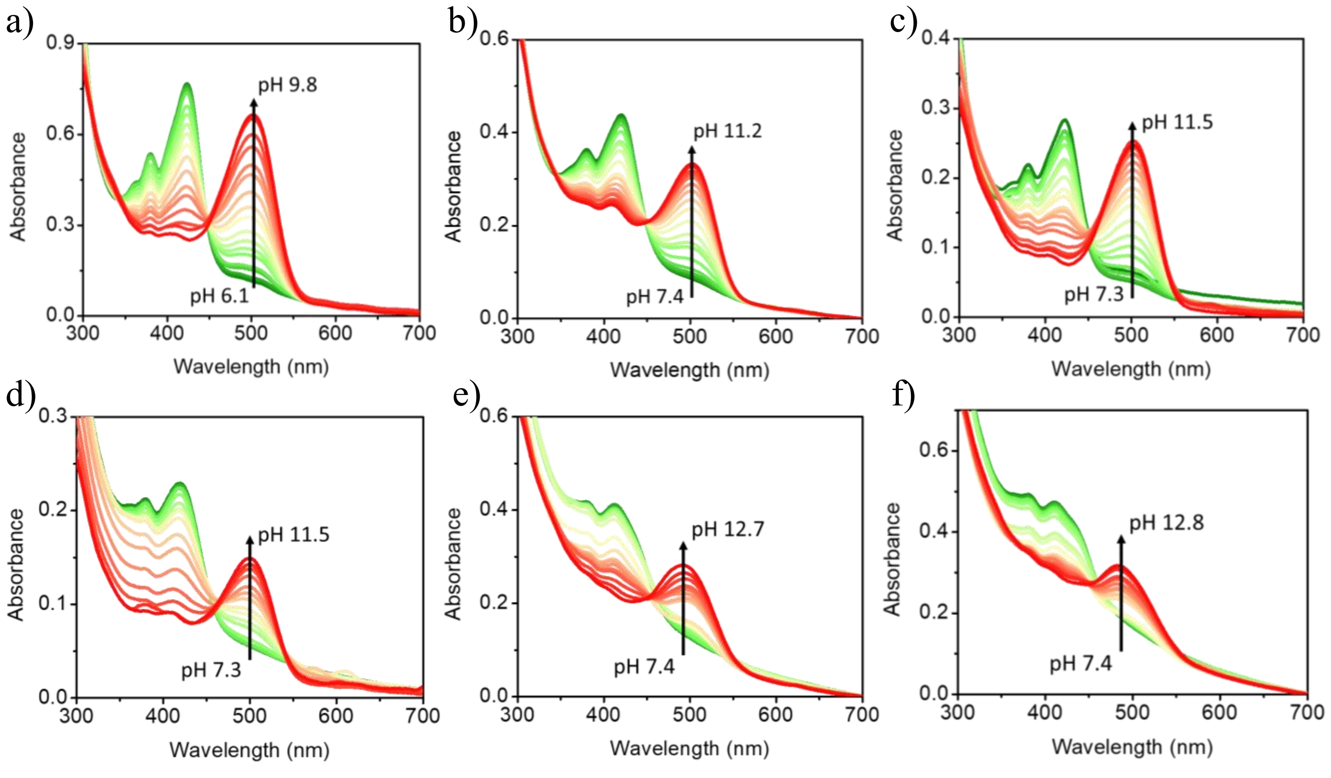


**Figure S2:** pH titration experiments carried out using UV-visible absorption spectroscopy showing the absorption of probe at different pHs for (a) DMPC, (b) DMPC:DMPA 9:1, (c) DMPC:DMPA 3:1, (d) DMPC:DMPA 1:1, (e) DMPC:DMPA 1:3, and (f) DMPA.

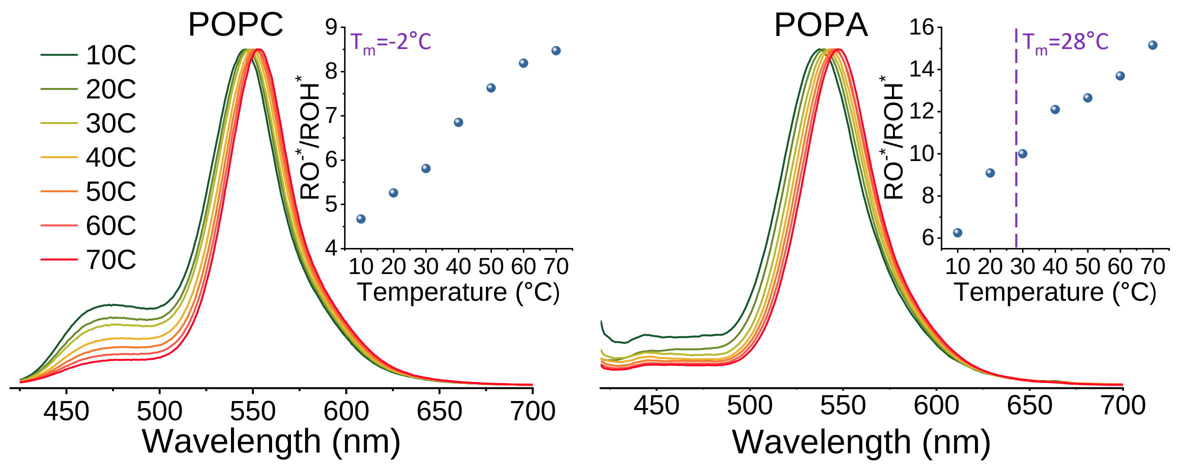


**Figure S3:** Temperature-dependent **s**teady-state emission spectra of $C_{12}$-HPTS inside POPC (right) and POPA (left). Insets showing the ratio of $RO^{-}*/ROH*$ calculated from the emission spectra vs temperature.

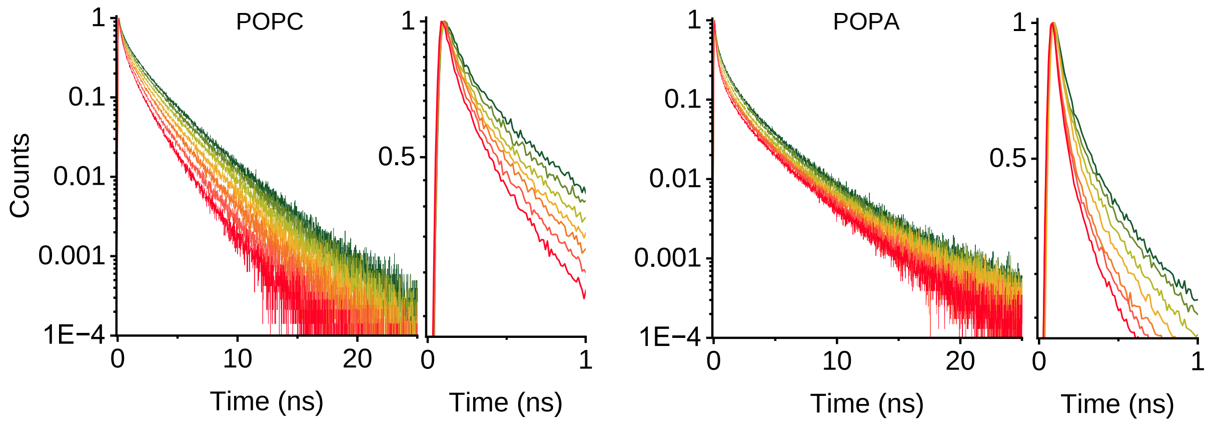


**Figure S4:** Time-resolved fluorescence showing the temperature-dependent behavior of $C_{12}$-HPTS inside POPC (right) and POPA (left). Insets showing the fluorescence decays for the first nanosecond.

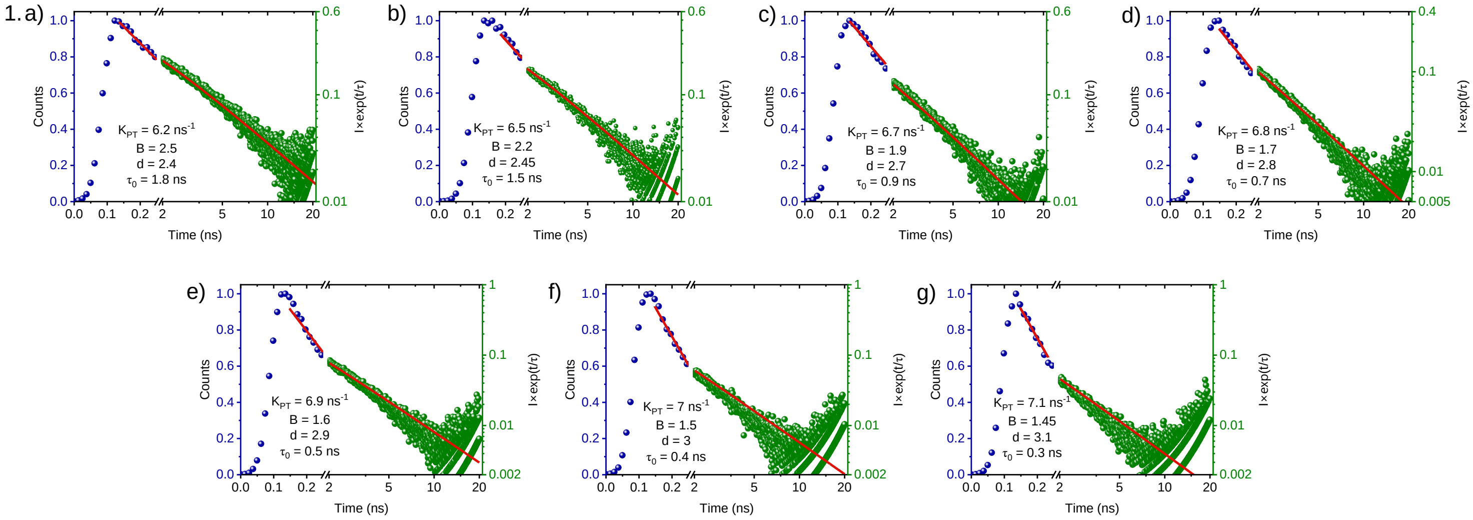
1. a)
Counts
Time (ns)
I×exp(t/τ)
$K_{PT}$ = 6.2 ns$^{-1}$
B = 2.5
d = 2.4
$\tau_0$ = 1.8 ns
b)
$K_{PT}$ = 6.5 ns$^{-1}$
B = 2.2
d = 2.45
$\tau_0$ = 1.5 ns
c)
$K_{PT}$ = 6.7 ns$^{-1}$
B = 1.9
d = 2.7
$\tau_0$ = 0.9 ns
d)
$K_{PT}$ = 6.8 ns$^{-1}$
B = 1.7
d = 2.8
$\tau_0$ = 0.7 ns
e)
$K_{PT}$ = 6.9 ns$^{-1}$
B = 1.6
d = 2.9
$\tau_0$ = 0.5 ns
f)
$K_{PT}$ = 7 ns$^{-1}$
B = 1.5
d = 3
$\tau_0$ = 0.4 ns
g)
$K_{PT}$ = 7.1 ns$^{-1}$
B = 1.45
d = 3.1
$\tau_0$ = 0.3 ns

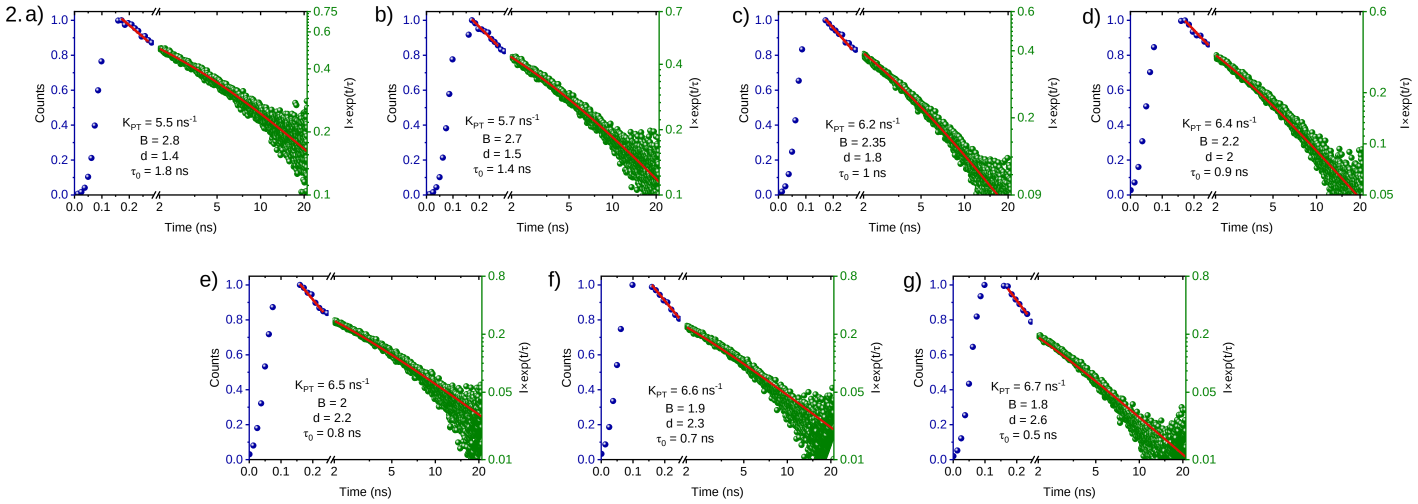
2. a)
Counts
Time (ns)
I×exp(t/τ)
$K_{PT}$ = 5.5 ns$^{-1}$
B = 2.8
d = 1.4
$\tau_0$ = 1.8 ns
b)
$K_{PT}$ = 5.7 ns$^{-1}$
B = 2.7
d = 1.5
$\tau_0$ = 1.4 ns
c)
$K_{PT}$ = 6.2 ns$^{-1}$
B = 2.35
d = 1.8
$\tau_0$ = 1 ns
d)
$K_{PT}$ = 6.4 ns$^{-1}$
B = 2.2
d = 2
$\tau_0$ = 0.9 ns
e)
$K_{PT}$ = 6.5 ns$^{-1}$
B = 2
d = 2.2
$\tau_0$ = 0.8 ns
f)
$K_{PT}$ = 6.6 ns$^{-1}$
B = 1.9
d = 2.3
$\tau_0$ = 0.7 ns
g)
$K_{PT}$ = 6.7 ns$^{-1}$
B = 1.8
d = 2.6
$\tau_0$ = 0.5 ns

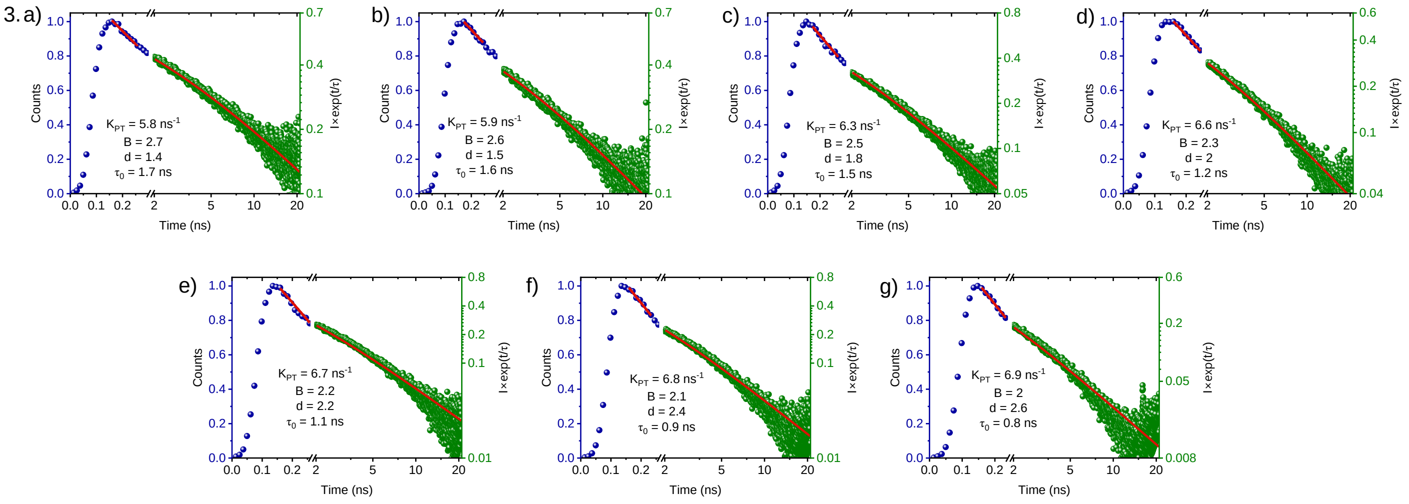
3. a)
Counts
Time (ns)
I×exp(t/τ)
$K_{PT}$ = 5.8 ns$^{-1}$
B = 2.7
d = 1.4
$\tau_0$ = 1.7 ns
b)
$K_{PT}$ = 5.9 ns$^{-1}$
B = 2.6
d = 1.5
$\tau_0$ = 1.6 ns
c)
$K_{PT}$ = 6.3 ns$^{-1}$
B = 2.5
d = 1.8
$\tau_0$ = 1.5 ns
d)
$K_{PT}$ = 6.6 ns$^{-1}$
B = 2.3
d = 2
$\tau_0$ = 1.2 ns
e)
$K_{PT}$ = 6.7 ns$^{-1}$
B = 2.2
d = 2.2
$\tau_0$ = 1.1 ns
f)
$K_{PT}$ = 6.8 ns$^{-1}$
B = 2.1
d = 2.4
$\tau_0$ = 0.9 ns
g)
$K_{PT}$ = 6.9 ns$^{-1}$
B = 2
d = 2.6
$\tau_0$ = 0.8 ns

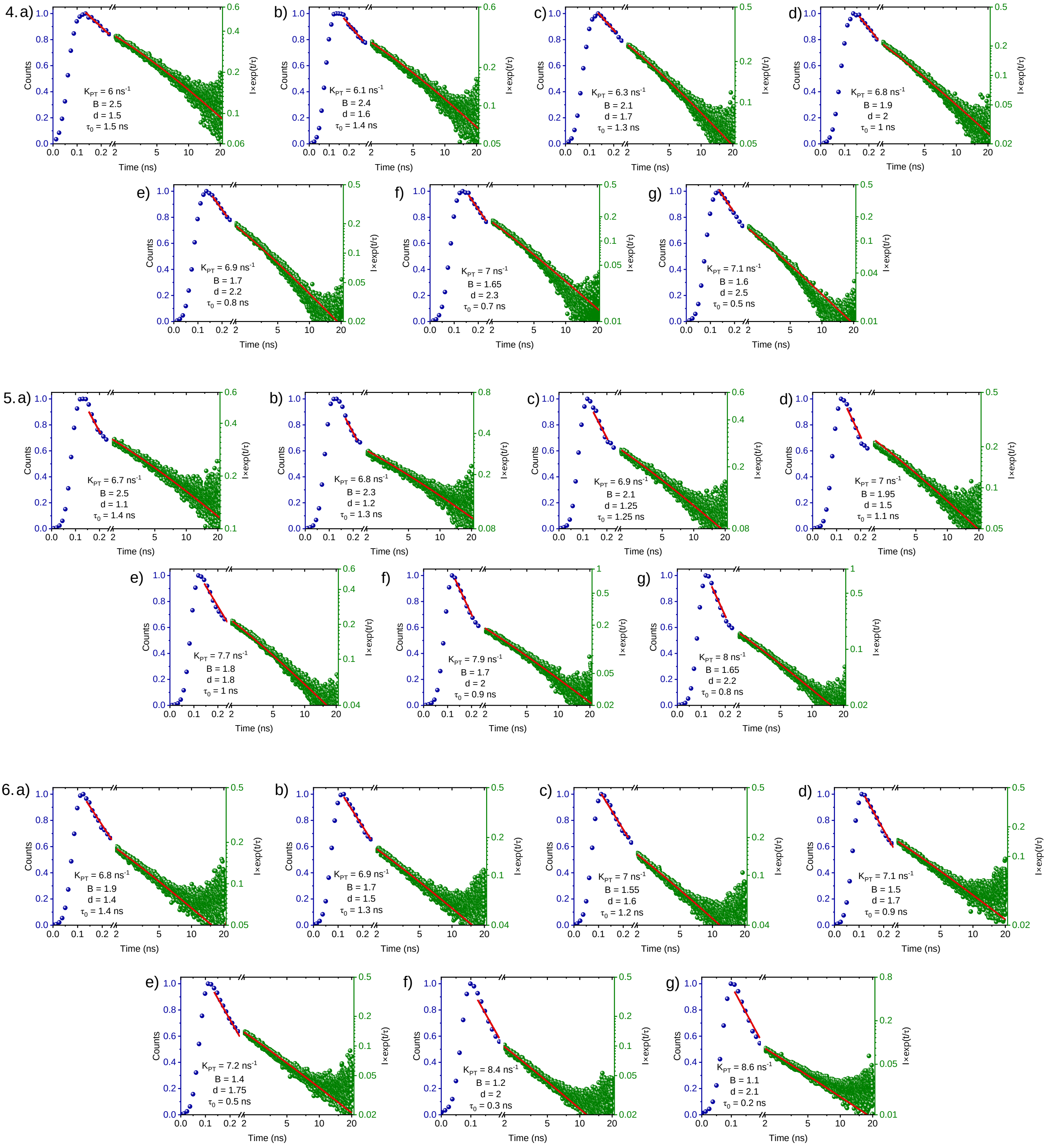


**Figure S5:** Fitting of all the fluorescence decays of (1) DMPC, (2) DMPC:DMPA 9:1, (3) DMPC:DMPA 3:1, (4) DMPC:DMPA 1:1, (5) DMPC:DMPA 1:3, and (6) DMPA with the theoretical model at (a) 10 ºC, (b) 20 ºC, (c) 30 ºC, (d) 40 ºC, (e) 50 ºC, (f) 60 ºC and (g) 70 ºC. The first few nanoseconds of data are fitted with equation (3) and latter nanoseconds are fitted based on equation (4) as shown in the main text.

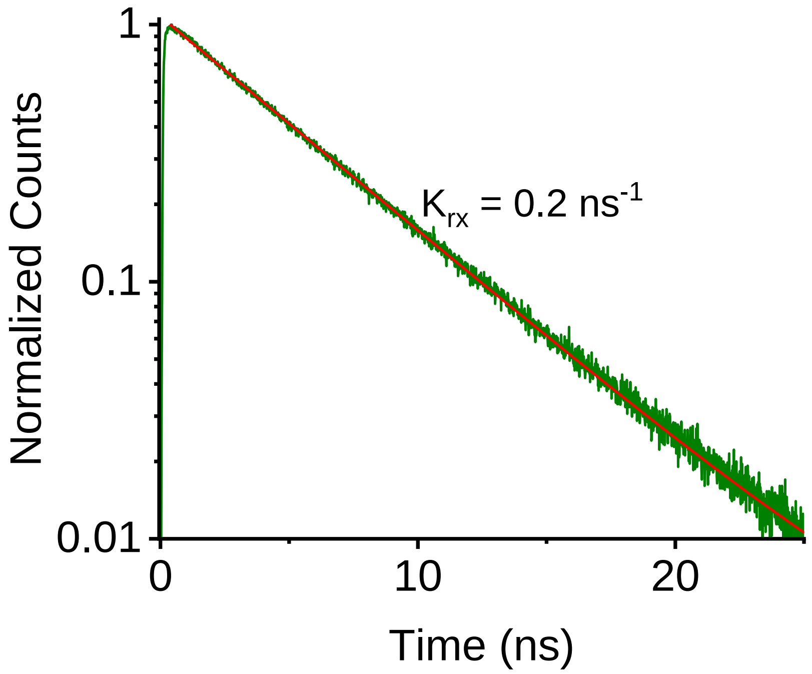


**Figure S6:** Exponential fitting of the decay of probe inside vesicles at 550 nm to find the rate constant.

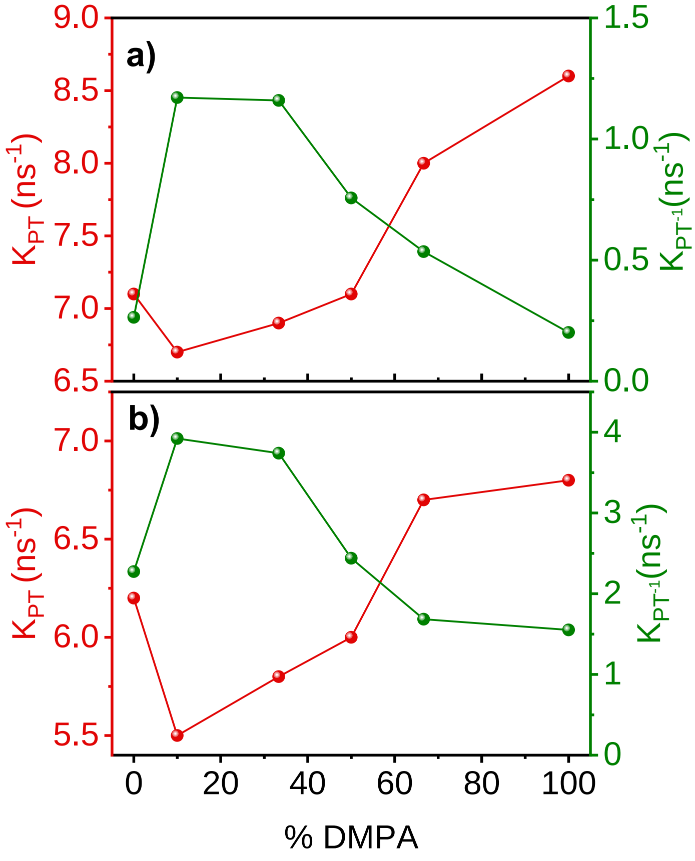


**Figure S7:** $k_{PT}$ and $k_{PT}^{-1}$ of all the mixture membranes at (a) 70 ºC and (b) 10ºC.

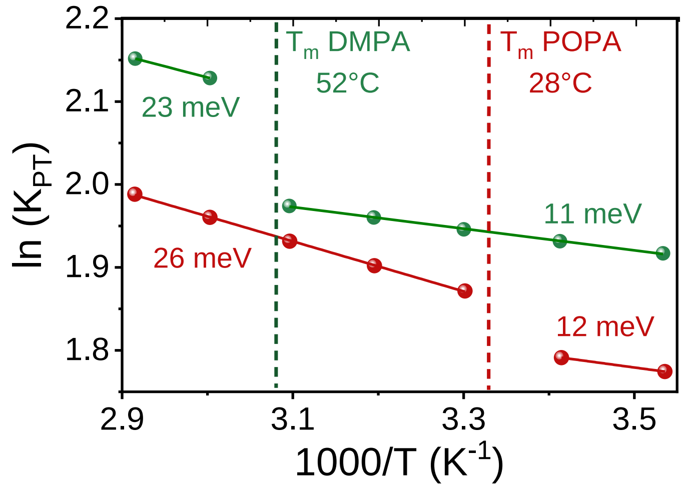


**Figure S8:** The change of $k_{PT}$ as a function of temperature for POPA (red) in comparison to DMPA (green).

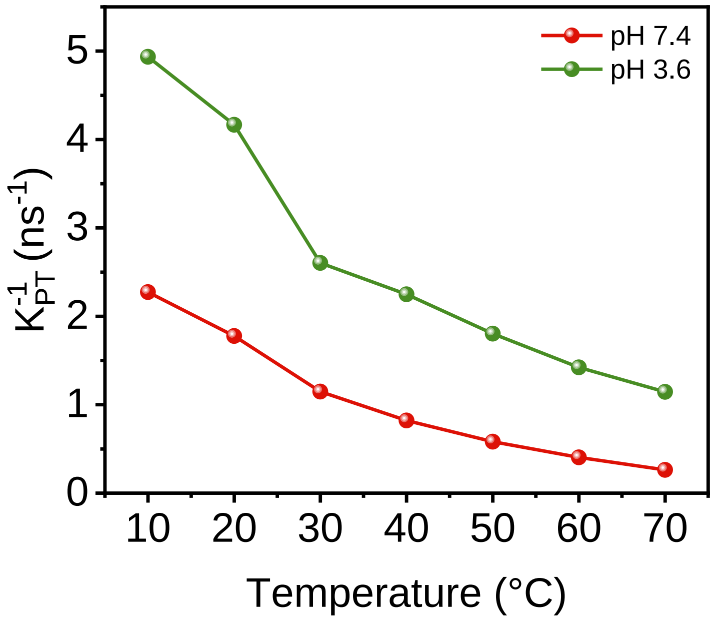


**Figure S9:** The temperature-dependent change in $k_{PT}^{-1}$ at two different pHs.

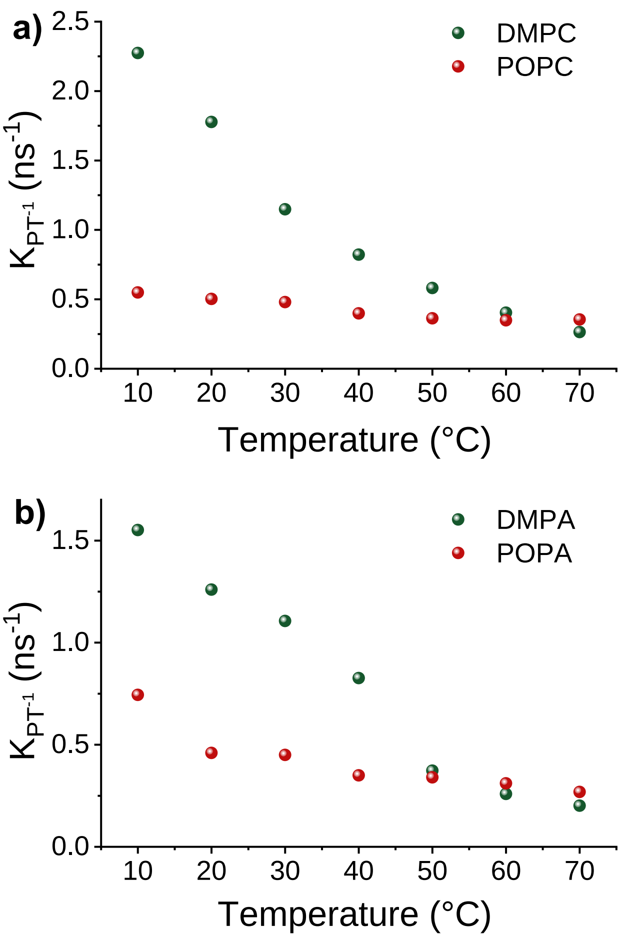


**Figure S10.** Comparison in the temperature-dependent change of the $k_{PT}^{-1}$ values between (a) DMPC and POPC, and (b) DMPA and POPA.

**MD Simulations:**

To explore the characteristics and dynamics of DMPA/DMPC bilayers, various membrane systems were generated using the CHARMM-GUI membrane builder (1). These systems comprised different mixtures of DMPC and DMPA at ratios of 1:0, 0.9:0.1, 0.75:0.25, 0.5:0.5, 0.25:0.75, and 0:1. The lipid bilayers were solvated using the TIP3P water model, maintaining 50 water molecules per lipid to ensure adequate solvation (2). Each PA/PC system contained 600 lipids, and Na+ ions were added to neutralize the system's total charge. Molecular dynamics (MD) simulations were performed using the GROMACS 2021 software suite (3) and the CHARMM36 force field (4).

Initially, energy minimization was performed on the membrane systems using a steepest descent algorithm. Subsequently, an NVT (constant Number of particles, Volume, and Temperature) ensemble was applied, with a leap-frog integrator and a 1 fs time step. In the NPT (constant Number of particles, Pressure, and Temperature) ensemble, a 2-fs time step was used. The simulations employed a Nose-Hoover thermostat across several temperatures: 283, 293, 305, 315, 323, and 343 K, with a time constant of $\tau_t = 2$ ps (5, 6). Pressure coupling was handled semi-isotropically using the Parrinello-Rahman barostat, set at 1 bar with $\tau_p = 5$ ps (7). During equilibration, positional and dihedral restraints were applied, with gradually decreasing force constants. Bond lengths were constrained using the LINCS algorithm (8).

During the production phase, the hydrogen mass repartitioning technique was employed, enabling a 4 fs time step (9). Van der Waals interactions were smoothly turned off between 1.0 and 1.2 nm through a force-based switching function (10), while long-range electrostatic interactions were calculated using the particle mesh Ewald (PME) method, with a grid spacing of 0.1 nm and a cutoff distance of 1.2 nm (11).

Area per Lipid and Membrane Thickness: A custom MATLAB script determined the membrane area per lipid (APL) and membrane thickness (MT). Each lipid molecule was described by the midpoint between P and C2 atoms' position. Such points were used for membrane thickness calculation; specifically, they were calculated as the difference between average height (z-axis) values of described points in opposite leaflets. This was followed by Voronoi tessellation to obtain the individual APL for each lipid molecule in every simulation time step. The APL dataset was histogrammed, and the APL value was obtained from the peak value of the fitted Gaussian function.

Bending Rigidity: The real-space fluctuation (RSF) method was used to determine the bending rigidity of investigated membrane systems (12). Specifically, for each lipid in each time step a splay was calculated, from which a distribution was calculated. Lipid splay is defined as the divergence of the angle formed by the directors of neighboring lipids providing that they are weakly correlated. The obtained distribution is fitted to obtain the bending rigidity coefficient.

Lateral Diffusion Coefficient: The lateral diffusion of lipid molecules was quantified using the Diffusion Coefficient Tool plugin (13). This coefficient is derived from Einstein's equation, utilizing the mean square displacement (MSD) of the selected molecular species. Specifically, the lateral diffusion for each lipid species was computed in the xy-plane, based on the positional data of phosphorus atoms. The analysis followed the procedure outlined in our previous study (14).

Interdigitation: The degree of interdigitation between the bilayer leaflets was assessed using the MEMBPLUGIN tool (15). This parameter represents the extent of mass overlap between opposing leaflets, quantified as the width of the overlap region within the bilayer.

Defects analysis: A protocol implemented by Boyd et al. was used for defect determination by acyl chain accessibility (17). Briefly, each bilayer atom (from lowest to highest) is described by its occupancy based on its Van der Waals radius. For head group atoms the occupancy is marked as polar, otherwise as a tail. The headgroup is defined as all atoms from top to down and including the C2 atom. This is followed putting square grids with a grid spacing of 0.5 Å over membrane. For each grid position, a line is led and is marked as either head or tail depending on the first collision with atoms' occupancy. As a result, both 2D map of acyl chain accessibility and a fraction of head-to-tail grid points is calculated. To ensure that observed acyl chain regions are not overestimated due to small gaps in head group coverage, the sphere probe analysis is performed. Specifically, for each point, a sphere probe of radius ranging from 0.1 nm up to 0.5 nm is set. If all the grid points in the range of this probe are classified as tails, those grid points are assigned as defects. In this way, only hydrophobic patches with given sizes and shapes are classified as true defects, which has the advantage over a previously determined 2D map of acyl chain accessibility.
